# Palynological studies on the selected members of *Fabaceae*

**DOI:** 10.1101/2024.04.02.587767

**Authors:** T.K. Pranav Raj, Shaju Ajin, B.S. Abhinna, Joseph Jojo

## Abstract

The investigation into pollen biology encompasses various stages such as pollen production, transfer to stigma, and the ensuing interactions leading to fertilization. Stigma plays a crucial role in providing an optimal environment for pollen germination and subsequent tube growth. However, conducting *in vivo* studies presents challenges due to the complexities of pistillate tissues, thus necessitating a reliance on *in vitro* cultures to comprehend pollen physiology. This study focuses on assessing pollen viability and germination in a selection of *Fabaceae* species, including *Canavalia gladiata, Centrosoma pubescens, Clitoria ternatea, Crotalaria pallida, Glyricidia maculata, Pueraria phaseoloides*, and *Vigna unguiculata*, utilizing tetrazolium testing and hanging drop culture methods. Results demonstrate a range of variability in pollen viability (from 19.38% to 80.78%) and germination (ranging from 23.90% to 79.81%), with *Crotalaria pallida* exhibiting the highest rates of viability and germination, whereas *Clitoria ternatea* and *Pueraria phaseoloides* displayed the lowest rates. This investigation contributes valuable insights into the pollen characteristics of *Fabaceae* species, hinting at possible connections between pollen viability/germination and seed/fruit set percentages.

## INTRODUCTION

Pollen grains are the haploid entities that embody the male partner in sexual reproduction of plants and pollen biology encompasses, the study of pollen production, its transfer to stigma and details of pollen pistil interaction leading to fertilization and seed set. The stigma provides an environment, which is most fit for pollen germination and pollen tube growth. However, pollen germination studies in vivo are not easily feasible because of the complications occurring in the pistillate tissue. The knowledge on the physiology and biochemistry of pollen germination and tube growth is therefore, mainly based on in vitro culture of pollen grains (Heslop-Harrison 1987; Steer and Steer 1989). Pollen viability is a critical aspect for any studies in pollen biology. It is imperative to know the extent of viability of pollen sample to be used for experimentation or pollination. Pollen viability can be determined by various methods like in vitro germination test, tetrazolium test etc. Among these tests, tetrazolium test based on the reduction of a colourless water-soluble tetrazolium salt to a reddish water insoluble substance formazone, in presence of dehydrogenase enzyme, is one of the simplest and most reliable tests. Pollen germination and tube growth are prerequisites for fertilization and seed set. Germination requirements of pollen vary appreciably from species to species. Apart from moisture, they generally require carbohydrate source, boron, calcium, hormones etc. Optimal conditions for pollen germination have been worked out only in few taxa. Brewbaker and Kwack (1963) have developed a medium suitable for pollen germination and tube growth of 86 species. At present there are several methods like hanging drop culture, sitting drop culture, suspension culture, surface culture etc. for culturing pollen. Of these methods, hanging drop method is the most commonly used one which essentially involves, suspending the pollen grains in a drop of nutrient medium on a cover glass hanging over the shallow depression of the cavity slide which is made airtight to prevent the evaporation of the nutrient medium. In this context, the present investigation is an attempt to study percentage of pollen viability using tetrazolium test and percentage of pollen germination by hanging drop method in selected members of *Fabaceae*.

Pollen viability refers to the ability of the pollen to perform its functions of delivering male gametes to the embryo sac. This functional property of pollen after their release from the anther varies greatly from species to species and its quality is assessed on the basis of its viability. The literature of the results of the experiments conducted to analyse the viability and germinability of pollen grains from certain plants are reviewed hereunder. In vitro pollen germination studies were conducted by Kumari et al. (2009) in *Trichosanthes dioea*. The results showed that the percentage of pollen germination was 70.20. Manonmani and Mekala (2016), investigated the pollen viability, germination and tube growth of some plant species in basic culture medium. The results of the pollen viability test revealed that the maximum staining was observed at 90 minutes at 55°C. The results of the germination and tube growth test revealed the medium containing growth hormones such as IAA and 2,4-D were the most suitable for rapid pollen germination and formation of long pollen tubes than the medium treated with sucrose, boron and KCI in all the experimental plant species.

## MATERIALS AND METHODS

During the present study, percentage of pollen viability and pollen germination of selected members of *Fabaceae*, such as *Canavalia gladiata, Centrosoma pubescens, Clitoria ternatea, Crotalaria pallida, Glyricidia maculata, Pueraria phaseoloides* and *Vigna unguiculata* were analyzed using tetrazolium test and hanging drop culture method respectively. Among the plants tested during the present study, *Canavalia gladiate* and *Vigna unguiculata* are cultivated as vegetables. The immature pods of these plants are used as vegetables. The flower buds of these plants were collected from different parts of Kozhikode district. Mature flower buds were collected by 10 am and were soon processed.

### TETRAZOLIUM TEST

Aqueous solution of 2, 3, 5, - tri phenyl tetrazolium chloride (0.5%) was prepared in 10% sucrose solution. Freshly prepared solution was used for all the experiments and the solution was kept in amber coloured bottle during the course of the experiment. One drop of this solution was placed on a clean micro slide and small amount of pollen grains were distributed uniformly in the drop. It was then covered by a cover slip. The slides were properly labelled and the preparation was incubated in a dark chamber at room temperature at relative humidity 95% for 60 minutes. After incubation period the slides were screened for red coloured and colourless pollen using a compound microscope and the values were recorded. The principle of the tetrazolium test is as follows. The viable pollen grains have the dehydrogenase group of enzymes which are absent in non-viable pollen. Dehydrogenase enzymes having strong oxidation / reduction potential, reduce the colourless water-soluble tetrazolium salt into water insoluble red coloured formazone giving red stain to the viable pollen grains (Martin et al., 2016).

### HANGING DROP CULTURE

In vitro pollen germination test is a simple experimental method for the study pollen germination, pollen tube growth and response of pollen system to physical and chemical factors. The composition of the germinating medium to obtain optimal response varies from species to species. Brewbaker and Kwack’s medium, Robert’s medium, Hodgkin and Lyon’s medium etc are the most commonly used germination media. Among this, the pollen culture media used in the present study was Brewbaker and Kwack’s (1963) medium, the composition of which is given in appendix I. A thin film of petroleum jelly was applied around the rim of the cavity of a cavity slide. Half a drop of culture medium was placed at the centre of a clean dry cover glass. Pollen grains were dusted into this medium by gently tapping the stamens. It was then mixed thoroughly with a needle to obtain a homogenous pollen suspension. The cover slip with the pollen suspension was inverted over the cavity of the cavity slide, such that the pollen culture drop was suspended in the centre of the cavity. It was made airtight by applying a gentle pressure over the cover slip. The slides were labelled properly. The cultures were incubated under room temperature for one hour with periodic observation. After the incubation period the pollen suspension was examined under a compound microscope. The slides bearing the pollen suspension were screened for germinated and non-germinated pollen grains and the values were properly tabulated (Jaiswal et al., 2019).

### EXPERIMENTAL PROTOCOL

In each of the species included in the present study, pollen grains from two different flower buds of were studied separately. Two slides were prepared for each of the species. A total of 500 pollen grains were scored from each of the species. The same protocol was followed for both pollen viability test and pollen germination test.

### ANALYSIS OF DATA

The results obtained during the present investigation were statistically analysed in terms of arithmetic mean and percentage.

## RESULTS

The results of the experiments conducted for the estimation of percentage of pollen viability and pollen germination in selected members of *Fabaceae* are as follows.

The percentage of pollen viability ranged between 19.38 and 80.78 with the highest value in *Crotalaria pallida* and lowest in *Clitoria ternatea* (Table No.1 & Graph). The percentage of pollen germination showed variation between 23.90 and 79.81 The percentage of pollen germination was maximum in *Crotalaria pallida* (79.81%) and minimum in *Pueraria phaseoloides* (Table No.2 & Graph).

**Table 01:** Table showing percentage of pollen viability in selected *Fabaceae* members.

| Sl No. | Name of the species | No. of pollen observed | No. of viable pollen grains | Percentage of pollen viability |
| --- | --- | --- | --- | --- |
| 1. | *Canavalia gladiata* | 531 | 169 | 31.82 |
| 2. | *Centrosoma pubescens* | 667 | 175 | 26.23 |
| 3. | *Clitoria ternatea* | 98 | 19 | 19.38 |
| 4. | *Crotalaria pallida* | 583 | 471 | 80.78 |
| 5. | *Glyricidia maculata* | 629 | 144 | 22.89 |
| 6. | *Pueraria phaseoloides* | 478 | 183 | 38.28 |
| 7. | *Vigna unguiculata* | 594 | 417 | 70.20 |

**Table 02:** Table showing percentage of pollen germination in selected *Fabaceae* members.

| Sl No. | Name of the species | No. of pollen observed | No. of germinated pollen grains | Percentage of pollen germination (%) |
| --- | --- | --- | --- | --- |
| 1. | *Canavalia gladiata* | 524 | 152 | 29.0 |
| 2. | *Centrosoma pubescens* | 517 | 244 | 47.17 |
| 3. | *Clitoria ternatea* | 502 | 131 | 26.09 |
| 4. | *Crotalaria pallida* | 659 | 526 | 79.81 |
| 5. | *Glyricidia maculata* | 471 | 202 | 48.79 |
| 6. | *Pueraria phaseoloides* | 502 | 120 | 23.90 |
| 7. | *Vigna unguiculata* | 508 | 328 | 64.56 |

**Graph 01:**
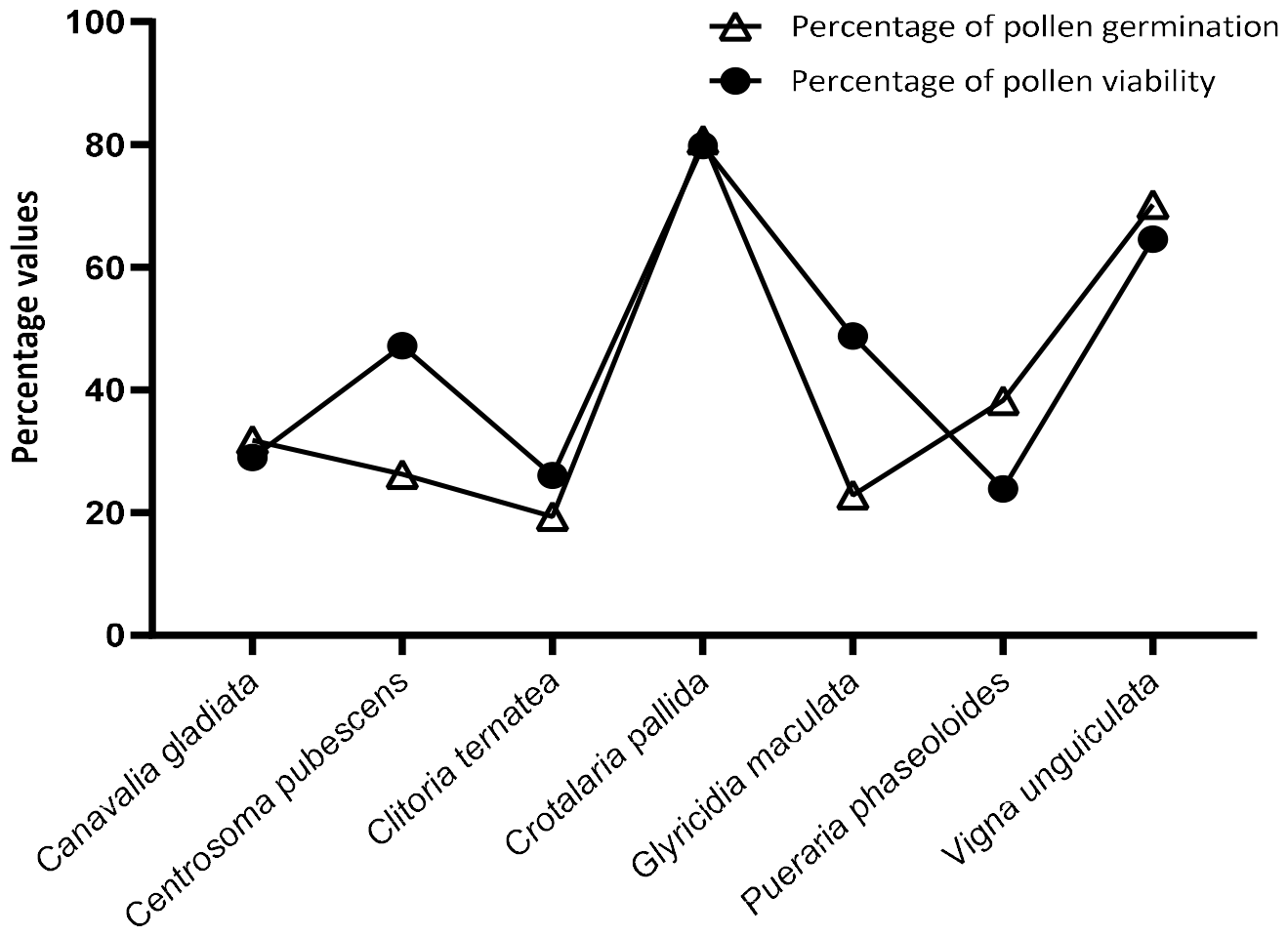
Graph showing the percentage of pollen viability and pollen germination in selected *Fabaceae* members

**Figure 01:**
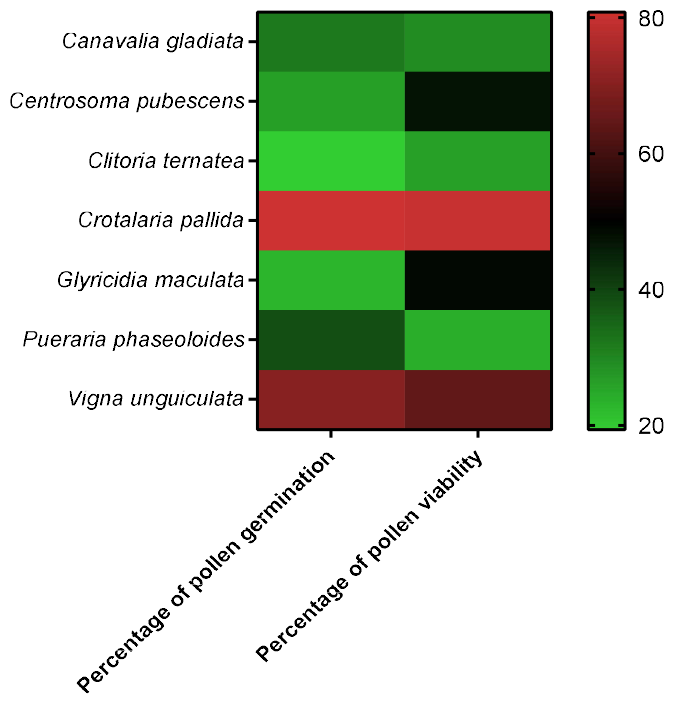
Heat map illustrating the percentage of pollen viability and pollen germination in selected *Fabaceae* members

## DISCUSSION

Germination is the first critical morphogenetic event in the pollen towards fulfilling its ultimate function of discharge of male gametes in the embryo sac. Because of the interference of pistillate tissue, the physiological and biochemical investigations of pollen germination and pollen tube growth in vivo are rather difficult. In vitro germination techniques have therefor been used extensively on a variety of pollen systems. Such studies have provided considerable information on the physiology and biochemistry of pollen germination and pollen tube growth (Shivanna and Johri, 1985; Shivanna and Rangaswamy, 1993). All the plants included in the present study showed pollen viability between 19.38% and 80.78% and pollen germination percentage between 23.90 and 79.81. This degree of pollen viability and germination may correspond to the degree of seed set in these members of *Fabaceae*. Visser (1955) showed a positive correlation between pollen viability and/or pollen germination and percentage of seed set. The plants included in the present study also showed high degree of seed set. This high setting percentage may be due to the high percentage of pollen viability and pollen germination showed by them.

## CONCLUSION

The present investigation, is an attempt to study the percentage of pollen viability and pollen germination of selected members of *Fabaceae*, such as, *Canavalia gladiata, Centrosoma pubescens, Clitorea ternatea, Crotalaria pallida, Glyricidia maculata, Pueraria phaseoloides* and *Vigna unguiculata* by tetrazolium test and hanging drop method in Brewbaker and Kwack’s (1963) medium. The percentage of pollen viability was maximum in *Crotalaria pallida* and was minimum in *Clitoria ternatea* Similarly the percentage of pollen germination was maximum in Crotalaria pallida and was minimum in *Pueraria phaseoloides*. The high percentage of pollen viability and pollen germination showed by these plants might be one of the reasons for the high percentage of seed set and fruit set showed by them.

## ACKNOWLEDGEMENT

This research was performed as a part of the dissertation, which was supported by Dept. of Botany, St. Joseph’s College (Autonomous), Devagiri, Medical College P O, Kozhikode, Kerala, India 673008

## AUTHOR CONTRIBUTION

JJ planned and designed the research. PRTK, AS, and ABS performed experiments, conducted fieldwork, analysed data, interpreted the data, and wrote the manuscript.

## DATA AVAILIABILITY

The authors confirm that the data supporting the findings of this study are available within article.

